# Single-Molecule Micromanipulation Studies of Methylated DNA

**DOI:** 10.1101/2020.07.29.227199

**Authors:** T. Zaichuk, J. F. Marko

## Abstract

Cytosine methylated at the 5-carbon position is the most widely studied reversible DNA modification. Prior findings indicate that methylation can alter mechanical properties. However, those findings were qualitative and sometimes contradictory, leaving many aspects unclear. By applying single-molecule magnetic force spectroscopy techniques allowing for direct manipulation and dynamic observation of DNA mechanics and mechanically driven strand separation, we investigated how CpG and non-CpG cytosine methylation affects DNA micromechanical properties. We quantitatively characterized DNA stiffness using persistence length measurements from force-extension curves in the nanoscale length regime and demonstrated that cytosine methylation results in increased DNA flexibility (i.e., decreased persistence length). In addition, we observed the preferential formation of plectonemes over unwound single-stranded “bubbles” of DNA, under physiologically relevant stretching forces and supercoiling densities. The stiffness and high structural stability of methylated DNA is likely to have significant consequences on the recruitment of proteins recognizing cytosine methylation and DNA packaging.

**Statement of Significance:** Despite countless structural and functional studies of DNA methylation, a key epigenetic mark in higher organisms, research towards the understanding of DNA intrinsic structural properties in the context of methylation layout representing different epigenetic landscapes is still in its initial stage. We utilize single molecule spectroscopy to analyze the effect of sparse symmetric and asymmetric 5-mC modification on the mechanical stability of long double-stranded DNA. Our findings establish that at physiologically relevant forces and supercoiling densities increased DNA flexibility of non-CpG methylated DNA translates to the high structural stability.

## Introduction

5-methyldeoxycytosine (5-mC) is a major DNA modification in higher eukaryotes and is the most studied epigenetic mark that occurs on ~ 3–8% of cytosines in mammals [1]. Since its discovery in 1948 [2] the 5-mC modification has been demonstrated to regulate a variety of biological processes including X chromosome inactivation [3], regulation of gene expression [4, 5], cell differentiation [6, 7] and chromatin compaction [8–10].

In metazoan genomes, DNA methylation has been traditionally considered to be restricted to symmetric modification of 5-mC at CpG loci. In bacterial and viral DNA, 5-mC functions within diverse sequence contexts and methylated CpG motifs are not frequent. This permits the vertebrate innate immune system to distinguish unmethylated CpG dinucleotides via the TLR9 pattern recognition receptor that initiates a signaling cascade triggering the interferon response and pathogen elimination [11].

The genomic landscape of CpG elements in humans has few distinct structural and functional patterns. Dispersed throughout the genome, sparsely methylated CpG dinucleotides limit the activity of transposable elements and maintain genomic stability. During tumorigenesis, such hypermethylation is progressively lost [12]. CpG methylation levels have bimodal distribution patterns in gene regulatory regions. Promoters with low CpG content are hypermethylated and are associated with tissue-specific genes. By contrast, promoters with densely clustered CpG islands (CGIs) are usually undermethylated and are associated with ubiquitously expressed genes. During positive clonal selection in the metastatic cancer process, such CGIs are more susceptible to hypermethylation and gene silencing [13].

Up to half of the transcriptionally inactive gene-poor genome regions exhibit a reduced average DNA-methylation level. Such lamina-associated partial methylation domains (PMDs), in combination with histone marks, demarcate more heterochromatic late DNA replication regions for a long-range epigenomic organization [14]. A recent study showed that PMD demethylation is a common feature of diverse cancer types [15]. Within a local CpG sequence context, preferential PMD hypomethylation increases with age and correlates with somatic mutation density and mitotic cell division [15].

Most human studies focused on 5-mC in the CpG context. However, the extensive methylation of human mitochondrial DNA in a non-CpG context has been reported recently [16]. The systematic non-CpG cytosine methylation with relatively low frequency is evidenced in stem and embryonic cells [17], neurons [18, 19], and cancer cells [20] where it enriches low CpG density regions and expands the proportion of genome regulated by cytosine modification [5].

Several studies addressed the impact of hydrophobic C5′-methyl group on DNA’s structure. Molecular dynamics simulations have indicated that the C5′-methyl group clash with the 5′ neighboring sugar causes an overall local increase of DNA flexibility [21]. Without affecting the hydrogen bonding properties of the bases, the C5′-methyl group increases the hydrophobic interactions involved in base stacking [22, 23], which is a major contributor to the dsDNA stability [24, 25]. Bulky C5′-methyl group directly inhibits the CG step overtwisting and indirectly, but to a comparable extent, torsionally modulates the stacking energy of adjacent base-pair steps [26].

One of the first observations of unusual thermodynamic stability conferred by 5-mC came from the analysis of bacteriophage XP-12 unique DNA with all cytosine completely replaced by 5-mC [27]. This substitution results in a more than 6°C increase in melting temperature [28]. Denaturing gradient gel electrophoresis was used to reveal that melting depends on both the number and location of methylated bases within a DNA fragment, as well as establishing that 5-mC at a non-CpG context is more efficient in stabilizing melting domains [29]. The effect of methylation is additive since methylation occurring at both CpG and CpT dinucleotides resulted in an even more stable DNA helix [29]. 5-mC located not only within but also up to three base pairs away from the A-tract causes alteration of DNA net curvature as measured by changes in electrophoretic mobility [30].

Methylation also affects dsDNA conformational transition from B to the left-handed Z form. Judged by circular dichroism and ultraviolet absorption, much weaker ionic strength is required for a methylated duplex of alternating deoxycytidines and deoxyguanosines to stabilize the Z form [31]. Such Z-DNA stabilization happens because methyl groups in the Z form are less exposed to the solvent than in the B form, as revealed by X-ray structures comparative analysis of the methylated and unmethylated CpG polymers [32].

Insights from these foundational studies inspired the question of whether the increased thermodynamic stability of the double-stranded methylated phage DNA, changes in DNA stability caused by methylation of one or two bases in a complex DNA sequence, and unusual optical properties of CpG polymers, affected DNA micromechanics, namely the bendability and ease of strand separation.

Persistence length (A), roughly 50 nm (150 bp) for unmodified double-stranded DNA in physiological buffer conditions, is the property that quantifies polymer stiffness, and corresponds to the average distance along which DNA maintains its direction before deviating from it. The definition of persistence length includes both flexibility and curvature [33] and it is important to highlight that different base pair steps show different susceptibility for helical deformations and deformations induced by slide or shift changes [34].

The persistence length of methylated DNA has been studied by traditional approaches including circularization assay [35], sedimentation velocities [31], optical methods, as well as by single-molecule techniques [36]. The conclusions drawn from these studies are divergent, possibly because of the indirect nature of these measurements combined with experimental and data analysis differences.

For example, selective cytosine methylation of DNA sequences with greater local flexibility resulted in a reduction in intrinsic stiffness of DNA as measured at the population level by cyclization kinetics experiments and Monte-Carlo simulation [37]; it is unclear if this observation remains true for sequences with increased or average intrinsic bendability. Conflicting with this stiffening-reduction observation, studies of short sequences that used circularization, AFM, and MD simulation approaches indicated that methylation of d(CpG) steps led to a lower tendency to bend and circularize [37], indicating that methylation was increasing stiffness. A significant increase in the persistence length of the surface-adsorbed methylated forms measured by AFM has also been reported by other groups [38–40], including observations of incremental changes in persistence length of DNA molecules with two different degrees of methylation [39, 40]. However, this decreased flexibility and changed contour length [40] of methylated DNA has been attributed to be, at least in part, a result of dehydration of the hydrophobic hypermethylated DNA on the substrate interface.

The mechanical response of long DNA under stretching forces and twisting torsions and held far from any surfaces has been investigated by optical trapping nanometry techniques [41]. By contrast with results for short oligos, hypermethylation of ~ 3.3 kb CpG-dense dsDNA was observed to lead to a reduction of the persistence length. A recent single-molecule spectroscopy study demonstrated that the persistence length of a 500-bp dsDNA increased by 1.5-fold when the GC content doubled from 30% to 64% and that this stiffening completely disappeared upon CpG methylation [42].

Altogether, prior findings indicate that DNA length, sequence context, and extent of methylation might influence the DNA bendability. Yet, the effects of varied patterns of DNA methylation across functionally distinct regions are still insufficiently explored. Many studies have addressed the changes in DNA nanomechanics of hypermethylated CpG islands that are densely clustered in promoters of ubiquitously expressed genes. However, nanomechanical responses of methylated sequences of tissue-specific promoters with low CpG content are less explored. Physical properties of asymmetrically methylated DNA that exist transiently during replication or maintained at regions flanking CTCF binding sites as a hemimethylation signature of progenitor cells are also understudied [42]. Additionally, to understand the role of methylation in chromatin architecture, partial methylation domains with a reduced average DNA-methylation level need to be analyzed.

In order to fill in these knowledge gaps, we designed a study to determine the effect of non-clustered cytosine methylation in the CpG and non-CpG context on the micromechanics behavior of long DNA. To obtain accurate force-extension curves profiling we used magnetic force spectroscopy which offered several advantages including constant-force mode of operation and submicron precision in monitoring the length changes of studied macromolecules at physiologically relevant stretching forces and supercoiling densities. We quantitatively characterized micromechanical properties of methylated DNA and found that even sparse cytosine methylation results in increased DNA flexibility and stability against strand separation that likely has consequences on gene regulatory landscape, and long-range epigenomic organization.

## Materials and Methods

### DNA tethers for magnetic tweezers experiments

To eliminate plasmid DNA modifications, we created a “bare” pFOS1-based construct via PCR amplification. PCR primer sequences are listed in Supplementary Table S1. The resulting 6062 bp DNA has 1376 (22.7%) of dC instances, with 330 of them in the context of dispersedly distributed CpGs. From the same plasmid, we created two labeled ~390 and 225 bp handles using PCR and used them as megaprimers for ligase-free synthesis of supercoilable tethers. The handles were synthesized by standard PCR with 10% biotin-dUTP or digoxygenin-dUTP included for labeling. The megaprimers were then used with the 6062 bp DNA as template to create supercoilable DNAs with multiple biotins and multiple digoxygenins at their ends. By replacing dCTP with 5-methyl-dCTP (New England BioLabs, Ipswich, MA) in the nucleotide mix, we optimized PCR amplification to achieve approximately 30% methylation. All constructs were column cleaned.

The bare (non-methylated) PCR constructs were also methylated enzymatically by CpG methyltransferase M.SssI (New England Biolabs, Ipswich, MA). To ensure complete methylation, the reaction was replenished with fresh S-Adenosyl-L-methionine and enzyme during a 3h incubation. After heat inactivation, the methylated DNA was purified using a DNA Cleanup kit (Sigma-Aldrich, St. Louis, MO).

The extent of CpG methylation was estimated by the selective BstUI restriction enzyme (New England Biolabs, Ipswich, MA) that does not cut targets (5’-CG^v^CG) containing 5-mC. Fig. 1A shows the locations of the BstUI sites in the region of pFOS1 used in our experiments.

**Figure 1:**
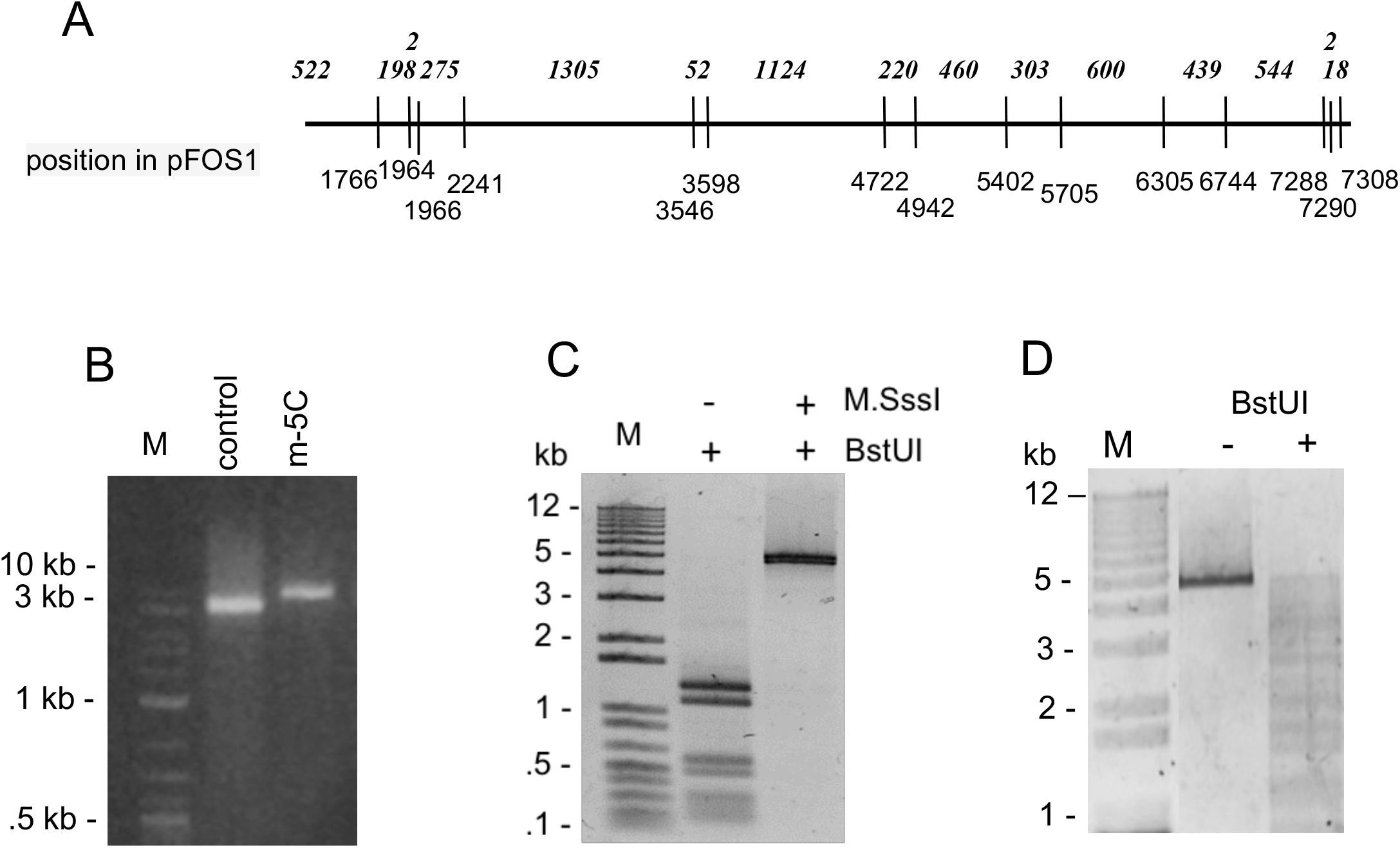
Verification of CpG methylation. (A). BstUI restriction sites and fragment sizes. (B). Control and methylated constructs created by PCR. The PCR products were run on 1% agarose gel and stained with ethidium bromide. (C) CpG methylation of ~ 6 kb PCR – created construct was carried out with M.SssI and the extent of methylation was checked by digestion with methylation-sensitive restriction enzyme BstUI. (D) The construct was randomly methylated by PCR and BstUI digestion produces a range of DNA fragments sizes.

### Magnetic tweezers experiment setup

Force-versus-extension curves were obtained by vertical magnetic tweezer measurements using a setup described in [43]. A simple microscope flow cell with a 40–50 μl volume was made using a #1.5 coverslip functionalized with 0.1 mg/ml anti-digoxigenin fragments (Sigma-Aldrich, St. Louis, MO). 0.5–1.0 ng/μl of DNA was added to the flow channel and incubated for 10 min. To form DNA tethers, 2.8 μm sized streptavidin-coated magnetic beads (50 μg/ml, Dynabeads MyOne Streptavidin, Invitrogen, Waltham, MA) were added and incubated for another 10 minutes.

Data was acquired using lab-written LabView software. By adjusting the distance between the magnet and the beads, we applied different stretching forces and monitored changes in the z position (or extension) comparative to a reference bead that reflects the end-to-end length of dsDNA. To ensure that only one DNA molecule is attached to the bead and to confirm that it is torsionally constrained, we applied positive or negative coils by rotating the magnets and tracked DNA extension for features indicative of plectoneme formation, at 0.3 pN and 1 pN. At low forces, the introduced supercoils induce the formation of plectonemes, contracting the DNA and results in symmetric “hat curves”, as seen when DNA extension is plotted against rotation. At 1 pN single DNA molecule extension decreases only when positively supercoiled. To minimize compaction and allow analysis within the low range of applied forces all data were acquired in 10 mM Tris, 1 mM EDTA, 100 mM NaCl (pH 7.5).

### Data analysis

The force-extension data with experimental extensions normalized to their full contour lengths were fitted to the classical wormlike chain (WLC) model of entropic elasticity to obtain the bending persistence length. For forces between 0.5 pN and 10 pN [44], it is accurately described by the equation *Z/L=1−(4AF/k*_*B*_*T)*^*1/2*^ where *Z/L* is the average extension scaled by the contour length, *F* is the tension along *z* direction, *k*_*B*_ is the Boltzmann constant, *T* is the absolute temperature and A is the effective persistence length. Using the expected linear relation between F^1/2^ and extension, the persistence length (A) values were extrapolated from plotting the reverse of the square root of force as a function of relative DNA extension normalized to the contour length. The model is specified as follows: 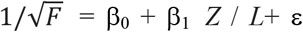 where β_0_, β_1_ are coefficients determined experimentally and ε is an error term. After having validated that the regression slopes were not significantly different across experiments, we combined all datasets for each construct into a single analysis and calculated persistence length values and errors [42, 45].

The length increase due to melting Δ*Z* was calculated from the extension at negative supercoiling density minus the extension at similar positive superhelical density, or Δ*Z* = 〈*Z*(−|σ|)〉 − 〈*Z*(+|σ|)〉. Superhelical density specifies a length independent value of supercoiling σ = ΔLk/Lk_0_ with ΔLk = Lk − Lk_0_ quantifies the extent of torsional deformation. For DNA composed of *N* base pairs, Lk_0_ = *N/γ* where γ is the helix repeat of 10.5 base pairs for a relaxed DNA double helix under physiological conditions. Linking number Lk = Tw+Wr, where twist Tw is the number of helical turns in the DNA and the writhe Wr is the number of times the double helix crosses over on itself forming plectonemic loops.

## Results

### Protection against BstUI digestion by incorporation of 5-mC

Three types of molecules were studied: the control sequence with 23% deoxycytidines lacked methylation altogether. This control sequence was enzymatically methylated by M.SssI [46] at the average frequency of one CpG element per 18 base pairs, providing a first methylated case for comparison with the control. Finally, using m-5C-PCR we prepared m-5C-saturated molecules, with methylation randomly distributed at an average rate of 1 per 15 bps within any context (CG, CHG, and CHH, where H indicates non-G nucleotides), providing a second methylated case for study.

Fig. 1B shows gel analysis the control and PCR-methylated molecules made using megaprimers, indicating that only a single product length has resulted, with a slight shift to lower mobility for the methylated molecule. Fig. 1C indicates the extent of methylation among the 15 BstUI CGCG sites for enzymatically and PCR-methylated constructs. Once enzymatically methylated at CpG sites, the DNA is completely resistant to BstUI cleavage, indicating near-complete methylation. By comparison, the 30%-methylated DNAs made by PCR showed only partial protection, producing BstUI cleavage products distributed as a smear from short lengths up to the length of the entire segments (Fig. 1D), instead of the precise ladder observed for the control construct (compare with Fig. 1C, middle lane).

### 5-mC decreases DNA persistence length

In order to determine the persistence length of our tethers, we probed the force-extension response of the three identical-sequence, PCR-created, ~6 kb DNA molecules with differing methylation, by measuring their extensions over a force range of approximately 0.3 to 1.2 pN. At this low to intermediate physiological force range, dsDNA elastic behavior is well described by the wormlike chain WLC model that takes into account local long-range flexibility. The effective persistence length for control and methylated DNA was determined by fitting the force-extension curve (Fig. 2). The least-squares fit of combined data for bare construct indicated an average slope corresponding to a persistence length of A = 44±2 nm, quantitatively close to previously reported results [33]. Coefficient estimates are found to be statistically significant at the 5% level, and additional details on coefficient values and significance values can be seen in Table S2.

**Figure 2:**
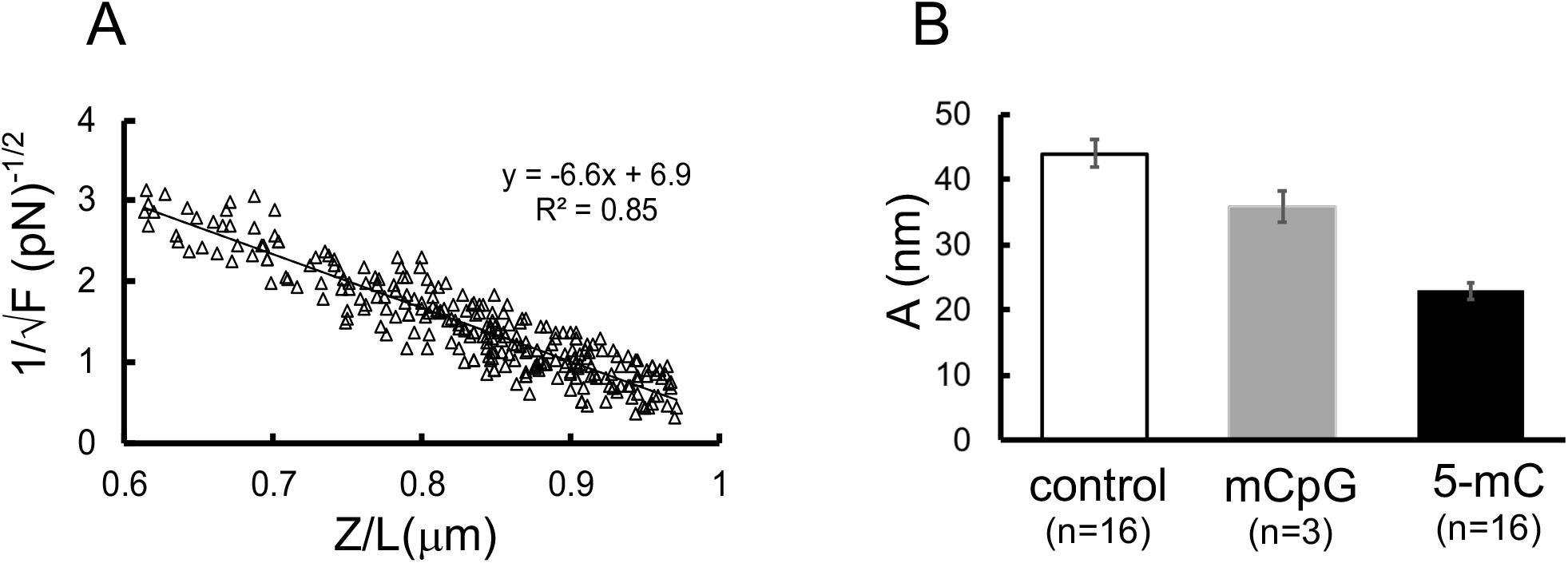
Persistence lengths of control, enzymatically methylated, and PCR-methylated DNA. (A). The persistence length value obtained from the force-extension curves of control DNA. The extension Z was measured at varying forces from 0.2 to 3 pN. To minimize noise, data is normalized to the contour length values measured at 5pN and relative elongation (z/L) values <0.97 were used for persistence length calculations. 1/√F is plotted as a function of z/L and persistence length A is determined from the slope of least-squares fit of data. The solid line represents the best fit of linear regression. (B). Comparisons of the persistence length values obtained from the force-extension curves of control (c, n=16), mCpG (n=3), and non-CpG (5-mC, n=16) methylated DNA; the number of experiments n for each construct is indicated.

For the enzymatically-methylated construct, a persistence length of A = 36±3 nm was measured. For the PCR-methylated construct, a persistence length of A = 23±2 nm was measured. Thus, both methylated versions of the DNA segment studied were found to be more flexible than the control (unmethylated) DNA.

### 5-mC increases DNA duplex stability against unwinding

We were interested to test whether 5-mC leads to a detectable perturbation in response of DNA to topological constraint over a range of physiologically relevant stretching forces and twisting torques. For torsionally constrained bare DNA under negative torque, there is a co-existence of twisted, plectonemic, and base-pair melted DNA [47–51] at forces between 0.5 and 1 pN [52] which provided us a method to make this test.

We determined the responses to unwinding and overwinding at a constant stretching force of 0.6 pN (referred to below as the “melting force” for bare DNA) that permits characterization of the transition from plectonemic to partially denaturated bubble-melted DNA. For bare DNA, as shown with open symbols and the dashed line in Fig. 3A, an increasingly positive number of turns at this melting force causes plectoneme formation and a decrease in DNA extension with a higher slope than that observed for negative turns. In the latter unwinding case, torque-melting of DNA leads to less plectoneme formation and a flatter extension response. The result is a highly asymmetrical extension response for over- and underwound bare DNA. This extension vs turns slope behavior is in agreement with dsDNA supercoiling behavior discussed in previous studies [49–52].

**Figure 3:**
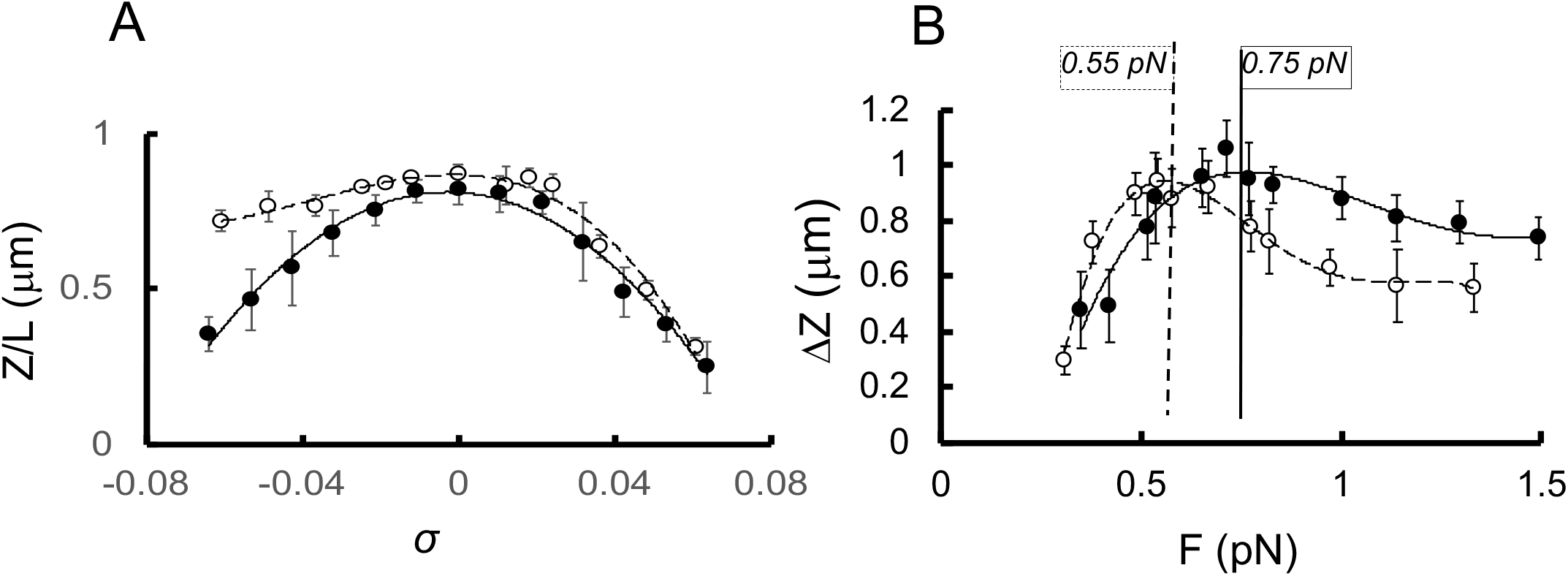
Extension - rotation and melting curves for single-DNA stretching and winding (+*σ*) / unwinding(−*σ*) measured using magnetic tweezers. **(A)** Rotation curves. Control (empty circles; dashed line) and methylated (filled circles; solid line) ~ 6 kb DNA constructs derived from pFOS1 by PCR (see Materials and Methods) held at constant force 0.6 pN at various superhelical densities *σ*. All data points are mean values of DNA extension that are collected within a 40s window and corresponding error bars are the standard deviation values of the DNA extension averaging. (B) Melting curves for control (empty circles; ; dashed line) and methylated (filled circles; solid line) DNA. Difference in extension (ΔZ= Z(−0.06)-Z(+0.06)) at the superhelical density *σ* +/−0.06 as a function of applied stretching force. Representative result of three independent experiments.

By contrast, for PCR-methylated DNA (Fig. 3A, solid symbols, solid curve) the “hat curve” is symmetric at this force; its extension is nearly the same underwound or overwound, indicating the absence of partial strand separation at 0.6 pN. In other words, methylated DNA is resistant to being unwound; i.e., requires more free energy to be strand-separated.

To further characterize the melting force for methylated DNA we studied the response of bare and PCR-methylated DNA that are over- and underwound, while being held under stretching forces ranging from 0.3 to 1.4 pN. The difference in extension (ΔZ) between underwound and overwound DNAs at superhelical density ±0.06 as a function of applied stretching force is shown in Figure 3B. Both DNA constructs show a ΔZ increase that correlated with an increase in applied force, and a slight decrease after reaching maximum. A higher applied force ~0.75pN was needed for methylated DNA to reach the maximum difference in extension, indicating a resistance unwinding-induced melting. This indicates that the 5-mC causes the well-resolved shift between coexisting dsDNA forms induced by negative supercoiling from partially melted towards plectonemic.

## Discussion

The results presented in this study describe the micromechanical features of long dsDNA methylated at 5-cytosines in the context of scattered CpG dinucleotides, or at any random context. Using force-extension measurements and the classical WLC model of entropic elasticity, we have demonstrated that enzymatic methylation at CpG dinucleotides decreases the persistence length of dsDNA from ~44 to 36 nm.

The most abundant motif in the PCR-methylated DNA used in this study is CHH, which is twice more frequent than CG or CHG sites. Although CpG sites are usually methylated on both DNA strands, this is not the case with non-CpG methylation because of sequence asymmetry [53] and even the methylation of symmetrical CHG motif departs significantly from a mirror-like complementary-strand pattern [54]. The persistence length of DNA with such a diverse methylation layout is ~23 nm. These data imply that increased DNA flexibility caused by 5-mC is context dependent and asymmetric methylation has a more profound effect on the bending persistence length. A similar trend in persistence length reduction by enzymatic and chemical dsDNA modifications (41.2nm and 39.1nm vs 44.6nm for control DNA) has been reported previously by Pongor et al. [41] for a ~3 kb hypermethylated dsDNA with high (59%) CpG content. However, our constructs have more pronounced changes in DNA flexibility. The lower extent of DNA methylation within sparse CpG context in our model or differences in techniques used, or a combination of both might contribute to such inconsistency. Further investigation also is necessary to elucidate the effect of structural particularities distinguishing between symmetrical and asymmetrical methylation on DNA flexibility.

Subsequent experiments to assess the effect of 5-mC on DNA topology were performed. It has been widely accepted that negative torsional strain occurring during transcription and replication causes interconversions between twisted, plectonemic, and melted DNA states that are related to DNA resilience and flexibility. DNA melting is a cooperative phenomenon that depends on the sequence length and composition, the number of modified bases, their position within a melting unit, and nearest-neighbor nucleotides [29, 52]. In our study, sparse random methylation of mostly non-CpG deoxycytidine increased the force (~0.75 pN vs ~0.6 pN for unmethylated counterpart) needed to separate methylated dsDNA strands and create denaturation bubbles during negative supercoiling at physiologically relevant superhelical density −0.06.

Opposite results were reported for short arbitrary sequences, studied by single molecule approaches [55]. The unzipping of short 20 bp dsDNA methylated duplexes requires a reduced force load and depends on the degree of methylation, as well as the localization of methylated sites along the chain. However, our results are in agreement with previously reported temperature-dependent melting experiments [28, 56-58] and DNA helix stability evaluation by denaturing gradient gel electrophoresis [29]. Moreover, it has been previously reported that hypermethylation of long CpG dense dsDNA disfavors strand unzipping and melting-bubble formation during mechanical relaxation after overstretching in the enthalpic regime of elasticity (10–60 pN) [41].

We review our current understanding of the 5-mC impact on DNA mechanical response. Using a magnetic tweezer, we confirmed that the effect of 5-mC modification can be detected by the changes in effective persistence lengths and described the effects of 5-mC on the stiffness and stability of sparsely methylated long DNA. The magnitude of the decrease in persistence length is more profound for methylation within the non-CpG context. More flexible DNA topology caused by 5-mC leads to a detectable shift to a higher force needed for the transition from plectonemic to melted DNA during negative supercoiling under physiological conditions.

There are multiple lines of evidence suggesting that the local DNA topology determines the selective usage of certain genomic regions as regulatory elements ([59] and refs therein). Data presented in this article provide a foundation for further study of how intrinsic DNA shapes and their nanomechanics might influence the organization of functionally distinct epigenetic landscapes. A key step will be to examine how the different DNA methylation profiles explored in this paper change binding properties of architectural and gene-regulatory proteins, histones/nucleosomes being a key example of the former, and transcription factors being a key example of the latter.

## Conclusion

Our findings confirm that non-CpG methylation increases DNA flexibility to a greater extent than methylation within CpG context. Furthermore, our results establish that decreased persistence length translates to the high structural stability of methylated DNA at physiologically relevant forces and supercoiling densities. The mechanistic insights from our study are likely to further our understanding of how DNA sequence context methylation contributes to defining the chromatin landscape and epigenetic deregulation.

## Data Availability

The authors declare that the data supporting the findings of this study are available within the paper and its Supplementary Information file. Additional raw data are available from the corresponding author upon reasonable request.

## Author Contributions

T.Z. conceptualized the project, conducted the experiments, analyzed the data, and wrote the original draft. J.M. developed methodology and provided instrumentation, oversaw the research activity, revised and edited the manuscript, and acquired the financial support for the project leading to this publication.

## Acknowledgements

We thank Sumitabha Brahmachari and Alexandra Lefevre for providing insightful suggestions for data analysis. This work is supported by NIH grants R01NS104041, R01-GM105847, U54-CA193419 (CR-PS-OC), and U54DK107980 (Center for 3D Structure and Physics of the Genome).

## References

[1] B.F. Vanyushin, S.G. Tkacheva, A.N. Belozersky. Rare bases in animal DNA, Nature 225 (1970) 948–9.

[2] R.D. Hotchkiss. The quantitative separation of purines, pyrimidines, and nucleosides by paper chromatography, J Biol Chem 175 (1948) 315–32.

[3] A.D. Riggs. X inactivation, differentiation, and DNA methylation, Cytogenet Cell Genet 14 (1975) 9–25.

[4] R. Holliday, J.E. Pugh. DNA modification mechanisms and gene activity during development, Science 187 (1975) 226.

[5] H. Han, C.C. Cortez, X. Yang, P.W. Nichols, P.A. Jones, G. Liang. DNA methylation directly silences genes with non-CpG island promoters and establishes a nucleosome occupied promoter, Hum Mol Genet 20 (2011) 4299–310.

[6] S.J. Compere, R.D. Palmiter. DNA methylation controls the inducibility of the mouse metallothionein-I gene lymphoid cells, Cell 25 (1981) 233–40.

[7] Z.D. Smith, A. Meissner. DNA methylation: roles in mammalian development, Nature Reviews Genetics 14 (2013) 204–20.

[8] I. Jimenez-Useche, D. Shim, J. Yu, C. Yuan. Unmethylated and methylated CpG dinucleotides distinctively regulate the physical properties of DNA, Biopolymers 101 (2014) 517–24.

[9] J.Y. Lee, J. Lee, H. Yue, T.-H. Lee. Dynamics of Nucleosome Assembly and Effects of DNA Methylation, Journal of Biological Chemistry 290 (2015) 4291–303.

[10] C.K. Collings, J.N. Anderson. Links between DNA methylation and nucleosome occupancy in the human genome, Epigenetics Chromatin 10 (2017) 18.

[11] A.M. Krieg. The role of CpG motifs in innate immunity, Current Opinion in Immunology 12 (2000) 35–43.

[12] M. Ehrlich. DNA methylation in cancer: too much, but also too little, Oncogene 21 (2002) 5400–13.

[13] A.R. Krebs, D. Schübeler. Tracking the evolution of cancer methylomes, Nat Genet 44 (2012) 1173–4.

[14] A. Salhab, K. Nordström, G. Gasparoni, K. Kattler, P. Ebert, F. Ramirez, et al. A comprehensive analysis of 195 DNA methylomes reveals shared and cell-specific features of partially methylated domains, Genome Biology 19 (2018) 150.

[15] W. Zhou, H.Q. Dinh, Z. Ramjan, D.J. Weisenberger, C.M. Nicolet, H. Shen, et al. DNA methylation loss in late-replicating domains is linked to mitotic cell division, Nat Genet 50 (2018) 591–602.

[16] V. Patil, C. Cuenin, F. Chung, J.R.R. Aguilera, N. Fernandez-Jimenez, I. Romero-Garmendia, et al. Human mitochondrial DNA is extensively methylated in a non-CpG context, Nucleic Acids Res 47 (2019) 10072–85.

[17] R. Lister, M. Pelizzola, R.H. Dowen, R.D. Hawkins, G. Hon, J. Tonti-Filippini, et al. Human DNA methylomes at base resolution show widespread epigenomic differences, Nature 462 (2009) 315–22.

[18] J.-H. Lee, S.-J. Park, K. Nakai. Differential landscape of non-CpG methylation in embryonic stem cells and neurons caused by DNMT3s, Scientific Reports 7 (2017) 11295.

[19] J.U. Guo, Y. Su, J.H. Shin, J. Shin, H. Li, B. Xie, et al. Distribution, recognition and regulation of non-CpG methylation in the adult mammalian brain, Nat Neurosci 17 (2014) 215–22.

[20] V. Patil, R.L. Ward, L.B. Hesson. The evidence for functional non-CpG methylation in mammalian cells, Epigenetics 9 (2014) 823–8.

[21] K. Liebl, M. Zacharias. How methyl–sugar interactions determine DNA structure and flexibility, Nucleic Acids Res 47 (2018) 1132–40.

[22] X. Teng, W. Hwang. Effect of Methylation on Local Mechanics and Hydration Structure of DNA, Biophysical Journal 114 (2018) 1791–803.

[23] G. Forde, L. Gorb, O. Shiskin, A. Flood, C. Hubbard, G. Hill, et al. Molecular structure and properties of protonated and methylated derivatives of cytosine, J Biomol Struct Dyn 20 (2003) 819–28.

[24] F. Kilchherr, C. Wachauf, B. Pelz, M. Rief, M. Zacharias, H. Dietz. Single-molecule dissection of stacking forces in DNA, Science 353 (2016) aaf5508.

[25] P. Yakovchuk, E. Protozanova, M.D. Frank-Kamenetskii. Base-stacking and base-pairing contributions into thermal stability of the DNA double helix, Nucleic Acids Res 34 (2006) 564–74.

[26] T.I. Yusufaly, Y. Li, W.K. Olson. 5-Methylation of Cytosine in CG:CG Base-Pair Steps: A Physicochemical Mechanism for the Epigenetic Control of DNA Nanomechanics, The Journal of Physical Chemistry B 117 (2013) 16436–42.

[27] T.-y. Feng, J. Tu, T.-T. Kuo. Characterization of Deoxycytidylate Methyltransferase in Xanthomonas oryzae Infected with Bacteriophage Xp12, European Journal of Biochemistry 87 (1978) 29–36.

[28] M. Ehrlich, K. Ehrlich, J.A. Mayo. Unusual properties of the DNA from Xanthomonas phage XP-12 in which 5-methylcytosine completely replaces cytosine, Biochim Biophys Acta 395 (1975) 109–19.

[29] M. Collins, R.M. Myers. Alterations in DNA helix stability due to base modifications can be evaluated using denaturing gradient gel electrophoresis, Journal of Molecular Biology 198 (1987) 737–44.

[30] Y. Hodges-Garcia, P.J. Hagerman. Cytosine methylation can induce local distortions in the structure of duplex DNA, Biochemistry 31 (1992) 7595–9.

[31] M. Behe, G. Felsenfeld. Effects of methylation on a synthetic polynucleotide: the B--Z transition in poly(dG-m5dC).poly(dG-m5dC), Proceedings of the National Academy of Sciences 78 (1981) 1619–23.

[32] S. Fujii, A.H. Wang, G. van der Marel, J.H. van Boom, A. Rich. Molecular structure of (m5 dC-dG)3: the role of the methyl group on 5-methyl cytosine in stabilizing Z-DNA, Nucleic Acids Res 10 (1982) 7879–92.

[33] M. Rittman, E. Gilroy, H. Koohy, A. Rodger, A. Richards. Is DNA a worm-like chain in Couette flow?: In search of persistence length, a critical review, Science Progress 92 (2009) 163–204.

[34] A. Pérez, F. Lankas, F.J. Luque, M. Orozco. Towards a molecular dynamics consensus view of B-DNA flexibility, Nucleic Acids Res 36 (2008) 2379–94.

[35] D. Shore, J. Langowski, R.L. Baldwin. DNA flexibility studied by covalent closure of short fragments into circles, Proc Natl Acad Sci U S A 78 (1981) 4833–7.

[36] J.P. Peters, L.J. Maher. DNA curvature and flexibility in vitro and in vivo, Quarterly Reviews of Biophysics 43 (2010) 23–63.

[37] A. Pérez, C.L. Castellazzi, F. Battistini, K. Collinet, O. Flores, O. Deniz, et al. Impact of methylation on the physical properties of DNA, Biophysical journal 102 (2012) 2140–8.

[38] P. Kaur, B. Plochberger, P. Costa, S.M. Cope, S.M. Vaiana, S. Lindsay. Hydrophobicity of methylated DNA as a possible mechanism for gene silencing, Phys Biol 9 (2012) 065001.

[39] V. Cassina, M. Manghi, D. Salerno, A. Tempestini, V. Iadarola, L. Nardo, et al. Effects of cytosine methylation on DNA morphology: An atomic force microscopy study, Biochimica et Biophysica Acta (BBA) - General Subjects 1860 (2016) 1–7.

[40] M. Wanunu, D. Cohen-Karni, R.R. Johnson, L. Fields, J. Benner, N. Peterman, et al. Discrimination of Methylcytosine from Hydroxymethylcytosine in DNA Molecules, Journal of the American Chemical Society 133 (2011) 486–92.

[41] C.I. Pongor, P. Bianco, G. Ferenczy, R. Kellermayer, M. Kellermayer. Optical Trapping Nanometry of Hypermethylated CPG-Island DNA, Biophysical Journal 112 (2017) 512–22.

[42] M.J. Shon, S.H. Rah, T.Y. Yoon. Submicrometer elasticity of double-stranded DNA revealed by precision force-extension measurements with magnetic tweezers, Sci Adv 5 (2019) eaav1697.

[43] D. Skoko, B. Wong, R.C. Johnson, J.F. Marko. Micromechanical Analysis of the Binding of DNA-Bending Proteins HMGB1, NHP6A, and HU Reveals Their Ability To Form Highly Stable DNA−Protein Complexes, Biochemistry 43 (2004) 13867–74.

[44] C. Bouchiat, M.D. Wang, J.F. Allemand, T. Strick, S.M. Block, V. Croquette. Estimating the Persistence Length of a Worm-Like Chain Molecule from Force-Extension Measurements, Biophysical Journal 76 (1999) 409–13.

[45] C.G. Baumann, S.B. Smith, V.A. Bloomfield, C. Bustamante. Ionic effects on the elasticity of single DNA molecules, Proc Natl Acad Sci U S A 94 (1997) 6185–90.

[46] M.V. Darii, N.A. Cherepanova, O.M. Subach, O.V. Kirsanova, T. Raskó, K. Ślaska-Kiss, et al. Mutational analysis of the CG recognizing DNA methyltransferase SssI: Insight into enzyme–DNA interactions, Biochimica et Biophysica Acta (BBA) - Proteins and Proteomics 1794 (2009) 1654–62.

[47] H. Meng, J. Bosman, T. van der Heijden, J. van Noort. Coexistence of twisted, plectonemic, and melted DNA in small topological domains, Biophysical journal 106 (2014) 1174–81.

[48] N. Gilbert, J. Allan. Supercoiling in DNA and chromatin, Current Opinion in Genetics & Development 25 (2014) 15–21.

[49] J.F. Marko. Torque and dynamics of linking number relaxation in stretched supercoiled DNA, Phys Rev E Stat Nonlin Soft Matter Phys 76 (2007) 021926.

[50] J.F. Marko, S. Neukirch. Global force-torque phase diagram for the DNA double helix: structural transitions, triple points, and collapsed plectonemes, Phys Rev E Stat Nonlin Soft Matter Phys 88 (2013) 062722.

[51] T.R. Strick, J.F. Allemand, D. Bensimon, V. Croquette. Behavior of supercoiled DNA, Biophys J 74 (1998) 2016–28.

[52] R. Vlijm, J. v.d. Torre, C. Dekker. Counterintuitive DNA Sequence Dependence in Supercoiling-Induced DNA Melting, PLOS ONE 10 (2015) e0141576.

[53] K. Shirane, H. Toh, H. Kobayashi, F. Miura, H. Chiba, T. Ito, et al. Mouse oocyte methylomes at base resolution reveal genome-wide accumulation of non-CpG methylation and role of DNA methyltransferases, PLoS Genet 9 (2013) e1003439.

[54] I. Ahmed, A. Sarazin, C. Bowler, V. Colot, H. Quesneville. Genome-wide evidence for local DNA methylation spreading from small RNA-targeted sequences in Arabidopsis, Nucleic Acids Res 39 (2011) 6919–31.

[55] P.M. Severin, X. Zou, H.E. Gaub, K. Schulten. Cytosine methylation alters DNA mechanical properties, Nucleic Acids Res 39 (2011) 8740–51.

[56] W. Szer, D. Shugar. The structure of poly-5-methylcytidylic acid and its twin-stranded complex with poly-inosinic acid, J Mol Biol 17 (1966) 174–87.

[57] J.E. Gill, J.A. Mazrimas, C.C. Bishop. Physical studies on synthetic DANs containing 5-methylcytosine, Biochimica et Biophysica Acta (BBA) - Nucleic Acids and Protein Synthesis 335 (1974) 330–48.

[58] L. Nardo, M. Lamperti, D. Salerno, V. Cassina, N. Missana, M. Bondani, et al. Effects of non-CpG site methylation on DNA thermal stability: a fluorescence study, Nucleic Acids Res 43 (2015) 10722–33.

[59] A. Pataskar, W. Vanderlinden, J. Emmerig, A. Singh, J. Lipfert, V.K. Tiwari. Deciphering the Gene Regulatory Landscape Encoded in DNA Biophysical Features, iScience 21 (2019) 638–49.

